# RFOptimization: Guiding Design Optimization with All-Atom Structure Prediction

**DOI:** 10.64898/2026.09.04.749184

**Authors:** Odin Zhang, Jiaqi Wang, Tuscan Rock Thompson, Ziyi You, Zihao Song, Frank DiMaio, David Baker

**Author notes:** These authors contributed equally (Odin Zhang, Jiaqi Wang, Tuscan Rock Thompson, Ziyi You).

## Abstract

Biomolecular interactions, including protein–protein interactions, protein–nucleic acid recognition, and protein–small molecule binding, underlie a wide range of biological processes and therapeutic mechanisms. Although recent *de novo* design methods can generate candidate binders for diverse molecular targets, practical design campaigns remain limited by low filter-passing rates and model-specific biases that arise when designs are optimized against a single predictor. Here, we present RFOptimization (RFO), a training-free framework for all-atom biomolecular binder optimization. RFO formulates binder improvement as a residue-wise mutational search problem, sampling candidate substitutions alternately based on gradient-guided sequence optimization using all-atom structure prediction models and a cycling-based sequence redesign strategy that alternates structure generation with an orthogonal predictor and MPNN-based sequence design to improve the *in silico* success rate of RFdiffusion-generated binders within minutes of computation. To reduce overfitting to any individual structure model, candidate mutations are further evaluated with orthogonal AlphaFold3 metrics as final filters. We demonstrate the generality of RFO across diverse design settings, including classical protein binder design, ligand-binding biosensor design, cyclic peptide design, and active site-aware enzyme design.

## 1 Main

Computational protein design plays a central role in enabling the rational engineering of biomolecular interactions for applications ranging from therapeutics to synthetic biology. However, the low *in silico* success rates of current *de novo* design pipelines remain a major bottleneck[1–10]. Diffusion-based models such as RFdiffusion[1] typically require sampling tens of thousands of designs followed by stringent computational filtering to obtain only a few candidates suitable for experimental validation. A complementary family of hallucination-based approaches uses structure prediction models directly to optimize protein sequences against their predicted structures and confidence outputs. BindCraft[4], for example, backpropagates through AlphaFold2 to optimize binder sequences and interfaces. More recent approaches, including BoltzDesign1, HalluDesign, and Protein Hunter[11–13], extend related predictor-guided optimization protocols to all-atom models and iterative sequence–structure design. Although these approaches differ in how they explore design space, they are primarily used for from-scratch generation. Consequently, during the process, a large fraction of generated candidates lies near the decision boundary but is discarded due to strict filtering criteria, despite containing promising structural features. Therefore, methods that refine these mid-tier designs within local sequence space would substantially improve the efficiency of design pipelines by increasing the fraction of generated candidates that are suitable for experimental validation.

We reasoned that AF3-like structure prediction models could be used to guide local optimization of existing designs. This presents two challenges. First, AF3-like models use a diffusion module for structure generation, which denoises a random coordinate cloud over many time steps. This diffusion-based structure module complicates direct backpropagation from structure-based objectives through each time step to the input sequence. Second, optimization against only a single structure prediction model may favor model-specific or adversarial solutions that improve its internal performance without generalizing to independent oracles. We set out to develop an optimization approach that overcomes these two problems.

## 2 Overview of RFOptimization architecture

RFOptimization is a training-free and modular framework for all-atom protein optimization that interleaves two complementary mutation strategies: a gradient-based mutation move and a cycling-based mutation move (Fig. 1). The underlying structure prediction models are modular and can, in principle, be replaced by other compatible AF3-like models; in our implementation, we use RoseTTAFold3 (RF3)[14] for gradient-guided optimization and Boltz[15, 16] for structure cycling. At each optimization step, RFOptimization selects between the two mutation moves with equal probability. The gradient-based move uses gradients from the optimization objective to guide residue mutations through RF3[14], whereas the cycling-based move combines Boltz structure prediction[15, 16] with ProteinMPNN[17] or LigandMPNN[18]-based inverse folding to resample sequences consistent with the predicted structure. These complementary moves combine objective-directed local search with structure-conditioned sequence exploration while reducing reliance on a single model. RFOptimization further provides a customizable objective function that can combine model confidence, distogram-derived structural criteria, and atom-level geometric constraints[14, 19, 20], enabling the same optimization framework to be adapted to diverse protein design tasks.

**Figure 1:**
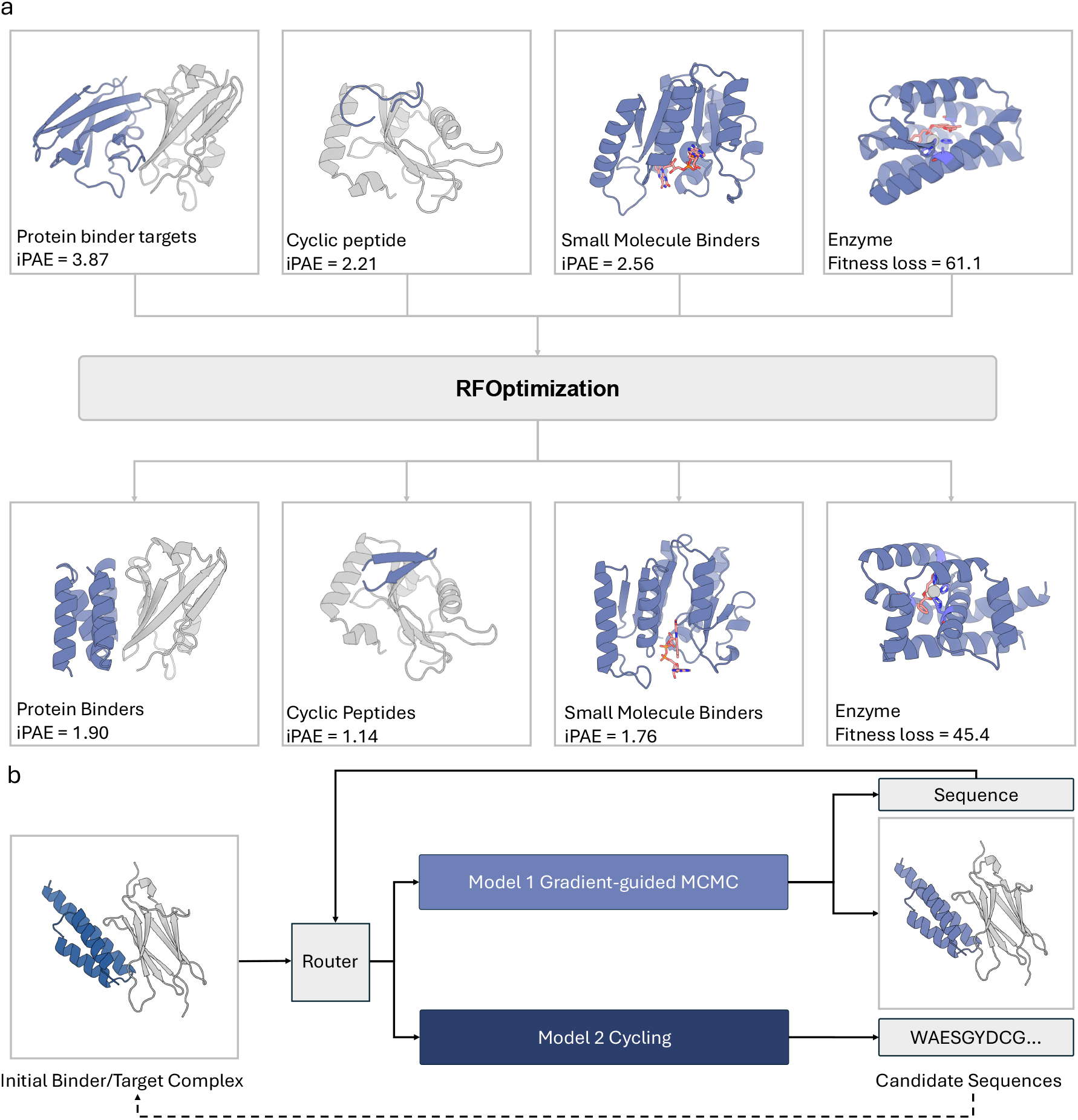
All-atom binder sequence optimization with RFOptimization. **a**, RFOptimization refines existing candidate designs across diverse molecular design tasks rather than generating structures *de novo*. Representative starting designs (top) and their corresponding optimized designs (bottom) are shown for protein binders, cyclic peptides, small-molecule binders, and enzymes. **b**, Hybrid optimization protocol. At each iteration, the method alternates between an RF3 gradient-guided MCMC step that backpropagates user-defined objectives to guide sequence mutations, and an AF3 structure conditioning step that applies partial diffusion to rectify the backbone followed by ProteinMPNN inverse folding to generate candidate sequences.

### 2.1 Gradient-based mutation move

A key challenge in gradient-based optimization of structure prediction models is that direct backpropagation through the diffusion module of RF3 and other AF3-like models is computationally prohibitive[14]. To address this issue, we apply a stop-gradient that blocks gradient flow through the diffusion module, bypassing it in the backward pass, and instead compute our optimization objectives from the differentiable heads attached to the Pairformer trunk, namely the distogram head and the confidence head[14, 19]. This design preserves our ability to express objectives tied to structural geometry, such as atom-level distance constraints between specific atoms or hydrogen bonding geometries, by formulating them as differentiable surrogates on the distogram and confidence outputs, so that the backward path from objective to input sequence remains fully differentiable[20, 21]. Upon backpropagating the objective loss, gradients flow from the confidence and distogram heads through the Pairformer to update the input sequence representation (Fig. 2a). Each optimization iteration requires only a single forward pass through the complete RF3 pipeline, taking approximately 45–60 seconds on a single A100 GPU. To prevent the optimizer from exploiting artifacts in the model’s continuous sequence representation, we maintain the sequence as a discrete one-hot encoding at every iteration rather than allowing it to drift into a continuous relaxation. After selecting a proposed mutation, we update the one-hot sequence and re-run RF3’s standard featurization pipeline, which regenerates the atom-level input features from scratch so that atom types, connectivity, and reference coordinates all reflect the new residue identity[14]. This keeps each iteration consistent with the model’s training protocol and ensures gradients are computed with respect to a valid discrete sequence. Users can additionally specify a mask indicating which residues are fixed and which are free to mutate, allowing optimization to focus on designable positions while preserving the functional motifs of interest. At each iteration, we select the residue mutation that yields the largest improvement in the loss function.

**Figure 2:**
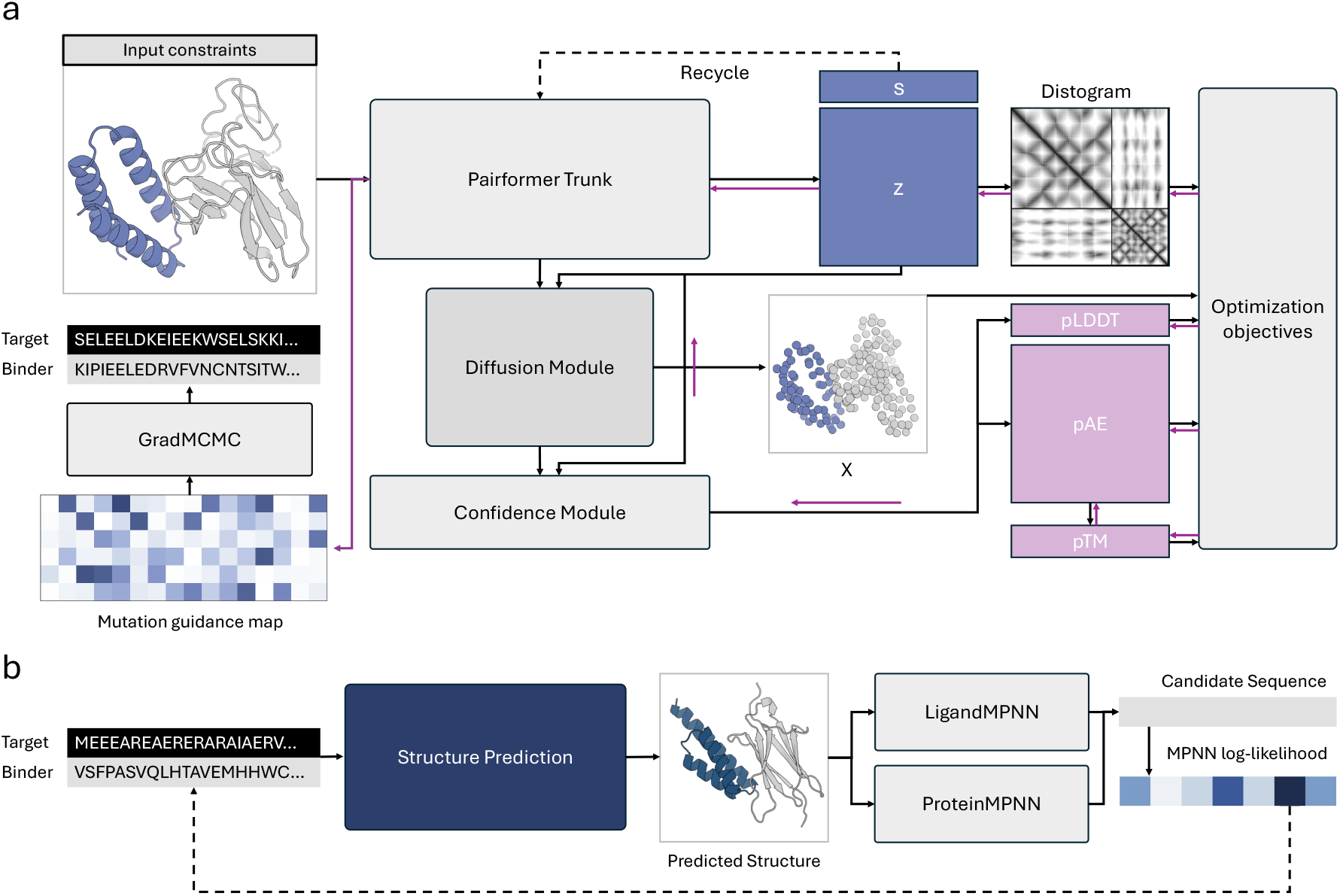
RFOptimization mutation proposal pipeline. **a**, Gradient-based mutation move. Black arrows indicate the forward prediction pathway and pink arrows indicate the direction of loss backpropagation. *s* represents the single representation and *z* represents the pair representation. User-defined optimization objectives, including pLDDT, pAE, and pTM, are computed from the confidence module and distogram heads. The resulting gradient attributions are used by gradient-guided MCMC to select sequence mutations, maintaining discrete one-hot encoding at every iteration. **b**, Cycling-based mutation move. The current binder and target sequences are passed to AF3-like models for structure prediction with partial diffusion to produce a predicted binder–target complex. The predicted structure is then inverse-folded using either ProteinMPNN for protein-only targets or LigandMPNN for targets containing small molecules or non-protein entities. Each method generates multiple candidate sequences conditioned on the predicted backbone (color strips represent per-residue sequence logits), and the highest-scoring candidate by MPNN log-likelihood is selected for the next optimization cycle.

Given the complex manifolds that describe binding interactions, we implement a gradient-guided Markov Chain Monte Carlo (MCMC) algorithm that combines the efficiency of gradients with the exploratory capability of stochastic sampling[22, 23]. The proposal is accepted according to the Metropolis–Hastings criterion where improvements will be accepted and deleterious mutations may be accepted with temperature-dependent probability, which allows the protein sequence to escape from local minima. To balance exploration and exploitation over the course of optimization, we implement geometric temperature annealing[24]. Early in optimization, we apply a high temperature that encourages broad exploration of sequence space, allowing the algorithm to deviate from poor initial states quickly to discover promising new regions. As optimization progresses, decreasing temperature focuses the search on local refinement to converge toward high-quality solutions. We find that this gradient-guided MCMC approach consistently achieves meaningful improvements within 1–10 iterations of optimization, which correspond to approximately 1–10 minutes of total runtime.

### 2.2 Cycling-based mutation move

The cycling step provides a complementary mutation strategy that proposes sequence changes through structure-consistent inverse folding rather than gradient-guided optimization. In this move, we first pass the current sequence into an external structure prediction model (Boltz in our implementation[15, 16]) to obtain a predicted all-atom structure of the binder–target complex. This prediction serves as an independent structural assessment using a different model than the RF3 backbone used for gradient computation, providing an orthogonal structural hypothesis for the current sequence. We then apply ProteinMPNN[17], or LigandMPNN[18] for small molecule and enzyme targets, to generate multiple candidate sequences that are consistent with the predicted backbone geometry. The highest-scoring candidate, as ranked by the MPNN sequence recovery score, is selected as the proposed sequence for the next iteration. During inverse folding, residues at the binding interface or within functional motifs can be fixed to preserve critical interactions, while the remaining designable positions are resampled.

### 2.3 Customizable objective function

The optimization objective function is highly customizable, allowing users to define differentiable combinations of structural and biophysical criteria for their needs. We organize the available objectives into three categories: confidence-based losses, distogram-based proxy losses, and atom-level geometric constraints. Confidence-based losses include interface predicted aligned error (iPAE), pLDDT scores, interface predicted TM-scores (iPTM), etc., that leverage RF3’s learned quality metrics to optimize predicted structure accuracy and binding[14, 19]. We also support several distogram-based proxy losses, including contact losses that encourage inter-chain contacts between binder and target or intra-chain contacts within the binder, and interface entropy losses that minimize the uncertainty of predicted distances at the binding interface. These proxy losses are computed directly from the distogram, which accelerates the optimization process. Together, the confidence and proxy losses approximate the energy landscape of protein binding, guiding the designs towards promising regions of sequence space.

In many protein design tasks, particularly in enzyme design[25–29], metal coordination, and covalent binder design[30], precise geometric control is a primary criterion that guides design. For these cases, we provide atom-level geometric constraint objectives that allow users to specify precise distance requirements between arbitrary atom pairs, which is helpful for enzyme active sites, metal coordination, or covalent interactions[14, 20, 21]. These three types of objectives enable a constrained search strategy through which the model can optimize complex atomic constellations.

In practice, design tasks often involve satisfying multiple and sometimes competing criteria, for example, optimizing confidences while simultaneously maintaining a catalytic geometry, and failure to satisfy any one criterion can lead to non-functional design. Existing generative approaches optimize a fixed, model-internal objective and offer limited control over which properties are prioritized, requiring users to rely on post hoc filtering to enforce additional criteria. RFOptimization addresses this by allowing users to construct composite objectives as weighted combinations of arbitrary design criteria, so that trade-offs between different requirements can be navigated directly during optimization rather than discovered after the fact. We also allow users to define their own custom criteria and implement new loss classes that compute differentiable metrics from RF3’s outputs. This “objective-oriented” paradigm represents a departure from the fixed fitness landscapes of generative models, where users directly specify what properties they want to optimize rather than retraining models for each design task.

## 3 *In silico* results

We first evaluate RFOptimization’s ability to improve designs generated by RFdiffusion[1]. These designs were optimized using RFOptimization with confidence losses as objective functions, and success was evaluated by refolding with AF3[19] to ensure that improvements reflect genuine structural quality rather than overfitting to model-specific minima in RF3’s[14] or Boltz’s[15, 16] confidence landscape. Across all benchmarks, we found that RFOptimization consistently transformed failed designs into candidates that pass the AF3-based structure prediction filters within 1–10 optimization iterations, corresponding to 1–10 minutes of computation time on a single A100 GPU. In the following sections, we analyze the ability of RFOptimization to improve designs across five distinct biomolecular design challenges and present a cross-model debiasing analysis that quantifies the success rate under a stringent consensus criterion across AF3, RF3, and Boltz.

### 3.1 Optimization of protein-binding protein binders and cyclic peptides

We first applied RFOptimization to protein-based binders, spanning both protein–protein interfaces and cyclic peptides. For protein–protein binders, we selected thirteen therapeutic targets previously benchmarked in protein binder design studies[1, 8, 31–33], including Programmed Death-Ligand 1 (PD-L1), Insulin Receptor (InsulinR), and Interleukin-7 Receptor-*α* (IL-7R*α*). For each target, the initial borderline designs are generated using RFdiffusion[1] as the starting point to RFOptimization.

We benchmarked the optimization of these borderline designs using mean-iPAE as objective functions. During optimization, designs were optimized using RoseTTAFold3[14] with target MSAs and templates, and success was evaluated by AF3[19] refolding under the criterion that AF3’s predictions satisfy iPAE *<* 2.5 and iPTM *>* 0.8. The average success rate improved from 6.73% to 22.31% within 10 optimization cycles, and RFOptimization improves the success rate on most targets (Fig. 3a). The largest absolute gains were observed on PDGFR (17.5% *→*50.0%), PDL1 (10.0% *→*40.75%), FGFR2 (0% *→*28.0%), PD1 (0% *→*27.5%), and VirB8 (0%*→* 27.25%), with several targets including CD3d, FGFR2, PD1, TGFb, Tie2, and VirB8 starting from a 0% baseline.

**Figure 3:**
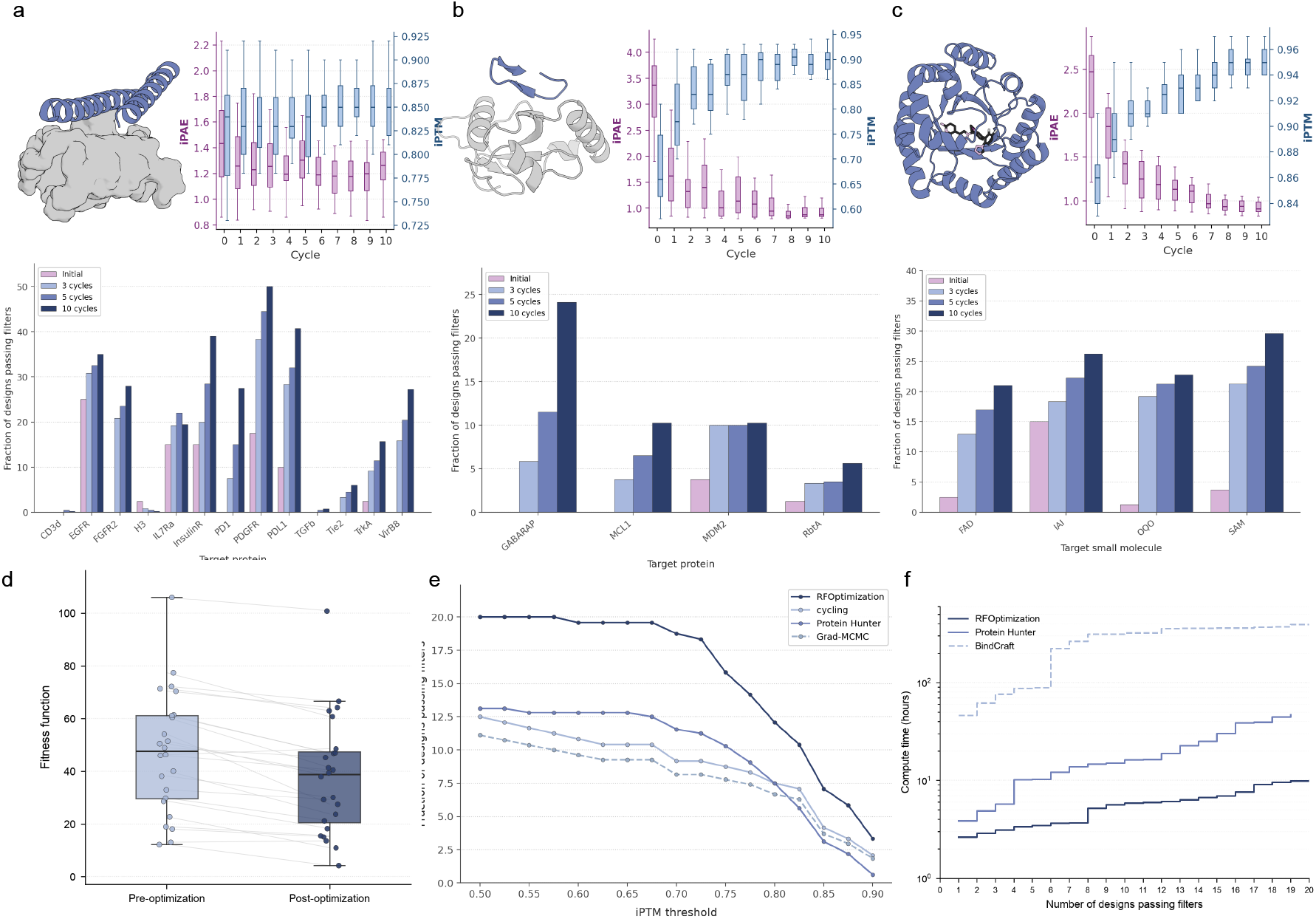
*In silico* benchmarking of RFOptimization across four biomolecular design challenges. In **a–c**, each block shows (left) an example optimized complex, (right) per-cycle distributions of iPAE (left axis) and iPTM (right axis) across 10 optimization cycles, and (bottom) the fraction of designs passing filters at successive checkpoints (initial, and best of 3, 5, and 10 cycles); success is defined by AF3 refolding under iPAE *<* 2.5 and iPTM *>* 0.8. **a**, Protein-binding protein binders across 13 therapeutic targets, improving the average success rate from 6.73% to 22.31%, with the largest gains on PDGFR (17.50%*→* 50.00%), PDL1 (10.00% *→*40.75%), FGFR2 (0% *→*28.00%), PD1 (0% *→*27.50%), and VirB8 (0% *→*27.25%). **b**, Cyclic peptides for four targets (GABARAP, MCL1, MDM2, RbtA), improving average success from 1.25% to 12.57%, with gains concentrated on GABARAP (0% *→*24.12%) and more modest improvements on MCL1 (0% *→*10.25%), MDM2 (3.75% *→*10.25%), and RbtA (1.25%*→* 5.63%). **c**, Small molecule binders for four ligands (FAD, IAI, OQO, SAM), improving average success from 5.62% to 24.91%, with SAM reaching 29.62%, IAI 26.25%, OQO 22.75%, and FAD 21.00%. **d**, Composite fitness function for enzyme active-site scaffolding (atom-level geometric constraints on the catalytic residues plus a substrate-pLDDT confidence term, where lower is better) before and after optimization across the 24 AME targets. **e**, Cross-model debiasing: fraction of designs passing a three-model consensus filter (iPAE *<* 2.5 and iPTM *> θ* under AF3, Boltz, and RF3 simultaneously) as the iPTM threshold *θ* is swept from 0.5 to 0.9, for full RFOptimization, the Boltz-cycling-only ablation (cycling), the gradient-only MCMC ablation with cycling disabled (Grad-MCMC), and Protein Hunter. RFOptimization passes at the highest rate across the entire sweep; at iPTM *>* 0.8 it reaches 12.08%, ahead of the cycling-only (7.50%) and gradient-only (6.67%) ablations and Protein Hunter. **f**, Compute efficiency on the 13 PPI targets, shown as cumulative GPU-hours to obtain a given number of designs passing the consensus filter for RFOptimization, Protein Hunter, and BindCraft. Amortized over full production cost, RFOptimization yields one filter-passing design per *∼*26 GPU-minutes, versus *∼*178 minutes for Protein Hunter and *∼*34 GPU-hours (*∼*2,030 minutes) for BindCraft.

We then extended the same approach to cyclic peptides, an important therapeutic modality that requires simultaneously satisfying binding-interface and cyclization-geometry constraints. We benchmarked RFOptimization on cyclic peptide design for four protein targets previously studied in *de novo* peptide design: GABARAP, MCL1, MDM2, and RbtA, and initial cyclic peptide scaffolds were generated using RFpeptides[34]. For cyclic peptide optimization, we applied mean-iPAE as the optimization objective, and the optimized designs were evaluated under the same AF3 refolding criterion: iPAE *<* 2.5 and iPTM *>* 0.8. RFOptimization improved the average success rate from 1.25% to 12.57% within 10 optimization cycles (Fig. 3b). The gains are concentrated on GABARAP (0%*→* 24.12%), with more modest but consistent improvements on MCL1 (0% *→*10.25%), MDM2 (3.75% *→*10.25%), and RbtA (1.25% *→*5.63%). The variation across targets suggests that RFOptimization’s effectiveness on cyclic peptide design depends substantially on the geometric tractability of the target binding pocket; GABARAP, with its well-defined hydrophobic LIR-binding groove, provides a more geometrically tractable surface for cyclic peptide optimization than MCL1, MDM2, and RbtA.

### 3.2 Optimization of ligand-binding protein binders

Ligand-binding proteins underpin a wide range of applications, from biosensing and molecular diagnostics to engineered light-harvesting systems[11, 30, 35–38], and present distinct challenges compared to protein–protein interfaces, as they require precise geometric complementarity to accommodate small molecule binding pockets. We selected four ligands previously studied in RFdiffusion benchmarks (FAD, IAI, OQO, and SAM)[2] and generated initial designs for each target. For optimization, we applied the same selection filter for borderline designs and used mean interface PAE as optimization objectives.

Optimized designs were refolded using RF3[14] with target MSAs and templated on target coordinates, and final success was evaluated by AF3[19] refolding under the criterion that iPAE *<* 2.5 and iPTM *>* 0.8. RFOptimization improved the average success rate from 5.62% to 24.91% across all four targets (Fig. 3c). FAD improves from 2.50% to 21.00%, OQO from 1.25% to 22.75%, SAM from 3.75% to 29.62%, and IAI from 15.00% to 26.25%. These gains are particularly notable because the initial success rates for several ligands were low, suggesting that many starting designs contained partially correct binding geometries that nevertheless failed the final AF3 confidence filter. With only three iterations of RFOptimization over the same RFdiffused starting structures, the success rate exceeds 10% on all four targets and exceeds 15% for IAI, OQO, and SAM. This indicates that a small number of local sequence refinements can substantially improve the predicted quality of protein–ligand complexes without requiring a new round of large-scale generation. The fact that RFOptimization can rescue designs across chemically diverse ligands, ranging from the large flavin cofactor FAD to the smaller methyl-donor SAM, suggests that RF3’s all-atom representation[14] provides a sufficiently general signal to refine pocket geometry for varied small molecule chemistries.

### 3.3 Optimization of enzymes and atomic motif scaffolding

*De novo* enzyme design and atomic motif scaffolding represent the most geometrically demanding protein design tasks, requiring precise three-dimensional positioning of specific atoms to catalyze a desired reaction[25– 29]. Prior work such as RFdiffusion2[3] has enabled scaffolding of catalytic motifs by jointly diffusing backbone frames and catalytic residue atoms, but performance degrades sharply on motifs split across many residue islands[14]. We evaluated RFOptimization on 24 targets from the atomic motif enzyme (AME) benchmark[3, 28] and generated the initial scaffolds per target with RFdiffusion[1] as optimization seeds.

For these targets we optimized a composite objective that combines atom-level geometric constraints on the catalytic residues with a pLDDT confidence term. The geometric constraints enforce the precise inter-atomic distances required for catalysis[25–29], while the pLDDT term simultaneously drives the surrounding scaffold toward a confidently folded structure[14, 19]; because RF3 operates on an explicit all-atom representation[14], the geometric component can specify distances between any arbitrary atom pairs. We track the value of this composite objective directly as a fitness function that scores how well a design jointly satisfies the catalytic geometry and confidence criteria, with lower values corresponding to tighter agreement with the target inter-atomic distances together with higher predicted substrate confidence. Comparing the fitness before and after optimization across the 24 AME targets, the distribution shifts downward: the median fitness decreases and 18 of the 24 targets improve from the start to the end of their optimization trajectory (Fig. 3d). This reduction is accompanied by a consistent increase in predicted substrate confidence across optimization cycles on all 24 targets, indicating that the optimizer moves designs toward more confidently predicted scaffolds while reducing the composite catalytic objective. This demonstrates the ability of RFOptimization to leverage RF3’s atomic-level weights to produce and refine complex catalytic geometries while maintaining global structural quality.

### 3.4 Ablation, cross-model generalization, and computational efficiency

We compared the performance of our two-branch protocol against simpler protocols that use only one of the two branches: gradient-based optimization, or structure-prediction/MPNN cycling. The full RFOptimization protocol interleaves a gradient-guided MCMC move through RF3 with a structure-prediction/MPNN cycling move through Boltz. The gradient-only ablation applies the RF3 gradient-guided MCMC step at every cycle with cycling disabled, and the cycling-only ablation applies only the Boltz cycling step. As an external baseline, we additionally included Protein Hunter[13], an independently developed cycling-based method. To test whether these protocols generalize across structure predictors rather than overfitting to the model used during optimization, we evaluated all methods under a three-model consensus criterion, counting a design as successful only if it simultaneously satisfied iPAE *<* 2.5 and iPTM *> θ* under RF3, Boltz, and AF3 (Fig. 3e). Across the entire threshold range, the full two-branch protocol achieved the highest consensus-filter pass rate. At the standard threshold of iPTM *>* 0.8, RFOptimization reached a consensus success rate of 12.08%, compared with 7.50% for the cycling-only ablation, 6.67% for the gradient-only ablation, and 7.50% for Protein Hunter. These results demonstrate that combining gradient-guided optimization with the cycling protocol improves cross-model consensus performance compared with either branch alone or a method confined to the confidence landscape of a single model.

We next examined whether these improvements generalize across structure prediction models. A central concern for optimization methods guided by structure prediction models is that apparent gains may reflect overfitting to the idiosyncrasies of the models’ confidence landscapes rather than improvements that transfer across predictors. Because RFOptimization derives its gradient-guided updates from RF3[14] and its cycling updates from Boltz[15, 16], optimized sequences could in principle exploit model-specific artifacts that inflate RF3 or Boltz confidence without producing corresponding improvements under other predictors. AF3[19] is never used during optimization and therefore serves as a held-out evaluation model. The advantage of RFOptimization under the three-model consensus criterion indicates that its improvements transfer across predictors rather than being confined to the confidence landscape of either optimization model.

Beyond improving model-agnostic design quality, RFOptimization also reduces the computational cost of obtaining candidates that pass the *in silico* filters. We compared the amortized GPU hours required to generate designs satisfying the consensus filter across the PPI benchmark (Fig. 3f). On a single A100 GPU, RFOptimization produced one filter-passing design per approximately 26 minutes, compared with approximately 178 minutes for Protein Hunter and approximately 34 hours for BindCraft[4]. The efficiency advantage reflects a shift from extensive sequence exploration to targeted local optimization, allowing RFOptimization to improve candidate designs within only a small number of optimization cycles.

## 4 Conclusion

RFOptimization optimizes biomolecular designs by inverting AF3-like models such as RF3[14] for gradient-guided sequence refinement and using structure cycling[15, 16] with a second structure prediction method to reduce adversarial solutions. By introducing gradient-guided search with MCMC sampling[22, 23], RFOptimization transforms borderline designs into higher-confidence candidates within minutes, addressing a critical bottleneck in current *de novo* design pipelines. Across four distinct design challenges, RFOptimization consistently improves AF3 refolding[19] success rates over the original RFdiffusion[1] outputs, with average improvements of 3.3*×* on protein–protein interfaces, 4.4*×* on small molecule binders, 10.1*×* on cyclic peptides, and 3.3*×* on atomic motif enzymes within only 10 optimization cycles.

Current limitations include dependence on the accuracy of the underlying structure prediction model and the quality of the initial designs provided as optimization seeds; as structure predictors improve[14–16, 19, 39–41], RFOptimization’s effectiveness should increase proportionally. Future work will focus on integrating RFOptimization with improved upstream sampling methods and large-scale design campaigns to further expand the accessible design space. Our *in silico* benchmarks across proteins, small molecules, cyclic peptides, and enzymes demonstrate the broad applicability of the method; RFOptimization provides scientists with a general platform for rapidly refining computationally designed binders toward any user-specified objective at atomic resolution.

## Acknowledgements

We thank Luki Goldschmidt and the IT team, Kandise VanWormer, Rafael Ticzon, and the lab management team for maintaining the computational and wet-lab resources at the Institute for Protein Design.

## Author Contributions

O.Z. led the architecture design and the overall study. J.W. contributed to model development and computational experiments. T.R.T. contributed to task design, base-model adaptation and evaluation, and loss-function design. Z.Y. contributed to the conceptualization of the study. Z.S. contributed to the enzyme design experiments. D.B. supervised the research and contributed to the overall study design.

## Supplementary Information

RFOptimization: Guiding Design Optimization with All-Atom Structure Prediction

### Supplementary Methods

## S1 Overview of RFOptimization

RFOptimization (RFO) is a training-free sequence-optimization framework for refining pre-existing biomolecular designs using pretrained all-atom structure-prediction models. Rather than training a task-specific surrogate model or generating a new binder from scratch, RFO treats a designed complex as an initial state and performs a local search over the sequence of a designated design chain. The search is guided by two complementary update mechanisms: (i) gradient-guided discrete sequence proposals derived from RoseTTAFold3 (RF3) [2], and (ii) structure-prediction/inverse-folding cycles that use an independently trained AF3-like predictor, Boltz [4, 5], followed by ProteinMPNN or LigandMPNN sequence redesign [6, 7]. The two update mechanisms are interleaved over a user-specified number of optimization cycles.

The key design principle is that the persistent optimization state is always a chemically valid, discrete biomolecular sequence. Gradients are used only to determine which amino-acid substitutions are locally favorable under an RF3-derived objective. RFO does not optimize a free continuous sequence embedding across iterations. Following every accepted mutation, the candidate sequence is converted back into a structure specification and passed through the RF3 featurization pipeline again, regenerating sequence- and atom-dependent features from the new residue identities.

For the binder-design experiments described in the main text, the first protein chain is treated as the designable chain and the remaining molecular entities provide the target context. The same implementation can accommodate protein targets, cyclic peptides, and non-protein entities represented through the all-atom input formats supported by the underlying predictors.

## S2 Notation and optimization state

Let a designable protein chain contain *L* amino-acid residues. Its discrete sequence at optimization step *t* is

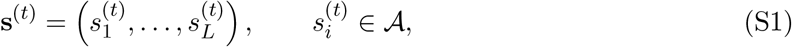

where *A* denotes the 20 canonical amino acids. The target sequence and any non-protein entities remain fixed unless explicitly designated otherwise.

RF3 represents residue identity using a categorical residue-type feature. During one differentiable RF3 evaluation, RFO clones this feature into a floating-point tensor

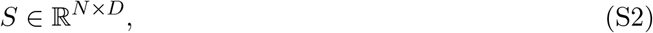

where *N* is the number of model tokens and *D* is the complete residue-type feature dimension used by the RF3/AtomWorks input representation. Gradient tracking is enabled for this temporary tensor. Mutation proposals are restricted to the first 20 canonical amino-acid channels.

The optimization objective is a scalar loss *ℒ*, and the local sequence-sensitivity matrix is

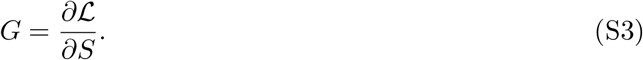

RF3 model parameters remain frozen throughout optimization; the derivative *G* is used only to construct candidate substitutions.

## S3 Input preparation and molecular representation

### S3.1 Initial complex

RFO accepts an initial biomolecular complex in PDB or mmCIF format. In the cycling implementation, chain A is treated as the designable binder by default. Protein target chains, ligands, and other molecular entities are retained as structural context.

If the input is supplied as mmCIF, the host-side cycling driver converts it to PDB for intermediate hand-off between model components. Predicted structures can subsequently be retained in either PDB or mmCIF format.

For protein chains, one-letter amino-acid sequences are reconstructed from standard residue names. Hetero residues and waters are excluded from protein-sequence extraction. Non-protein entities can be retained as CCD components or represented by SMILES strings through an optional AF3-style JSON template.

The AF3-style specification is particularly useful when information cannot be reconstructed unambiguously from a PDB representation, for example when a ligand should be specified by SMILES or when external sequence-search information should remain associated with a target chain.

### S3.2 Multiple-sequence alignments

Per-chain unpaired multiple-sequence alignments (MSAs) can be supplied as A3M files. When an MSA path is specified for a chain, the same chain-specific MSA information is propagated to both the RF3 and Boltz branches.

If no MSA is provided to Boltz, RFO explicitly requests single-sequence mode by setting the MSA field to

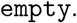

This avoids implicitly performing an external MSA search during the optimization loop.

### S3.3 Structural templates

The RF3 branch additionally supports fixed structural templates on a per-chain basis. This permits the target structure to be templated while the design chain changes sequence.

For binder-optimization experiments, target-chain templates and target MSAs can therefore be retained across all sequence iterations while only the binder sequence is modified.

### S3.4 Cyclic peptides

For cyclic-peptide optimization, chain A is explicitly marked as cyclic in the Boltz input representation. The remaining structure-prediction, mutation, and inverse-folding procedure is unchanged.

## S4 Sequence reconstruction and re-featurization after mutation

An amino-acid substitution changes more than the categorical identity of a residue. It can alter the expected atom set, atom masks, chemical connectivity, reference coordinates, and other sequence-dependent model features.

RFO therefore does not simply modify a residue-type tensor while reusing every other feature from the preceding sequence.

For each sequence evaluated by the gradient optimizer, the candidate sequence is projected back into a temporary CIF representation and subsequently passed through the RF3/AtomWorks transformation pipeline. The procedure can be written schematically as

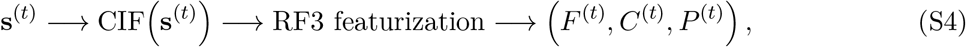

where *F* ^(*t*)^ denotes RF3 trunk features, *C*^(*t*)^ confidence-module features, and *P* ^(*t*)^ the remaining inference-pipeline outputs and molecular metadata.

The regenerated residue-type feature is cloned, converted to floating point, and assigned gradient tracking. Following a forward pass and objective evaluation, gradients are extracted with respect to this tensor.

This reconstruction step has two important consequences. First, every persistent optimization state corresponds to an actual discrete amino-acid sequence. Second, the atom-level inputs used by RF3 remain chemically consistent with the current residue identities.

## S5 RF3 gradient-guided optimization

### S5.1 RF3 inference during optimization

RFO loads a pretrained RF3 checkpoint in inference mode. Model parameters are not updated.

The public cycling configuration uses 10 RF3 recycles for the gradient branch and a diffusion batch size of one. The RF3 optimization wrapper supports both confidence-derived and distogram-derived objectives.

For distogram-only objectives, the wrapper can skip the structure-diffusion calculation and optimize directly from the differentiable distogram output. For confidence-derived objectives such as interface PAE or pLDDT, the full confidence-producing RF3 forward path is executed with gradient computation enabled.

Importantly, RFO does not directly optimize atomic coordinates. The derivative used by the sequence optimizer is always taken with respect to the temporary residue-type representation.

### S5.2 Gradient normalization

Raw sequence gradients can vary in scale as a function of sequence length, input composition, and objective. RFO therefore applies a global normalization before constructing mutation proposals.

For gradient matrix *G*, define the squared gradient magnitude associated with token *i* as

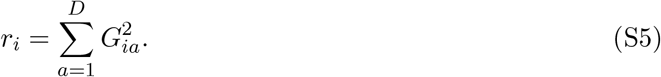

The effective number of positions receiving a non-zero gradient is

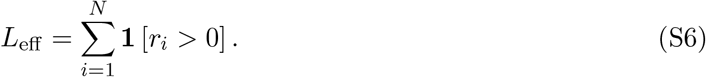

The normalized gradient used for proposal generation is

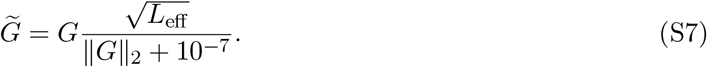

This transformation preserves the direction of the gradient while approximately controlling its global scale across sequences of different effective lengths.

### S5.3 Designable-residue mask

RFO constructs a Boolean mask over model tokens before proposing substitutions. Non-protein tokens are always excluded.

When binder-only optimization is enabled, the mask is intersected with the first-chain mask, such that only the binder sequence is mutable.

An optional interface-only mode further restricts the mutation space to residues located at a molecular interface. In this mode, pairwise atom distances are computed and an atom is classified as interfacial when it lies within

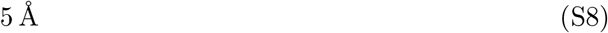

of an atom belonging to another chain. Interface atoms are mapped back to their corresponding residue tokens.

The user can additionally supply an explicit list of designable positions, allowing catalytic residues, binding motifs, covalent attachment points, or other functionally constrained sites to remain fixed.

### S5.4 Discrete gradient-derived mutation proposals

Because the optimization objective is minimized, candidate mutation logits are constructed from the negative normalized gradient,

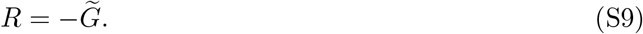

For deterministic mutation generation, the preferred amino acid at residue position *i* is

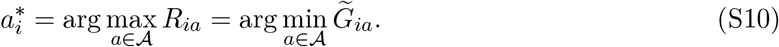

Only the 20 canonical amino-acid dimensions are considered.

A stochastic proposal mode is also implemented, in which the proposed amino acid is sampled according to

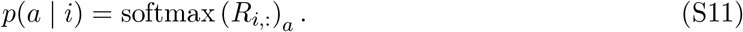

The standard GradMCMC configuration uses deterministic gradient-derived proposals.

For the first proposal, the logit associated with the amino acid already present at each residue is strongly penalized, forcing the optimizer to consider a non-identical amino acid.

The unconstrained gradient proposal can differ from the current sequence at multiple positions. If the number of changed positions exceeds the user-specified parameter `mutate_each`, RFO randomly selects `mutate_each` positions without replacement and retains only those substitutions.

The default cycling configuration sets

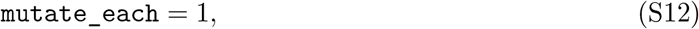

so that each individual GradMCMC proposal differs from the current accepted sequence by at most one amino acid.

## S6 Gradient-guided Markov chain Monte Carlo search

### S6.1 Current state and proposal evaluation

Let **s**^(*t*)^ denote the current accepted sequence with objective value

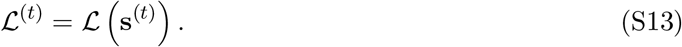

A candidate sequence **s***^′^*is generated from the current gradient. The proposed sequence is then completely re-featurized and independently evaluated by RF3 to obtain

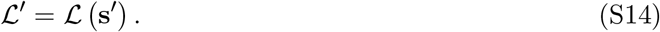

For a single RF3 checkpoint, define

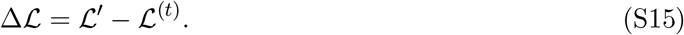

If Δ*ℒ <* 0, the candidate is always accepted. Otherwise, it is accepted with temperature-dependent probability

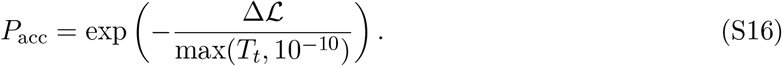

The sequence transition is therefore

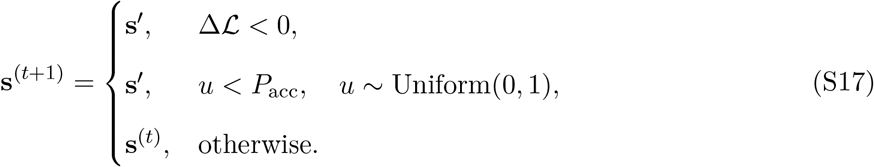

Because the implementation does not include an explicit proposal-density ratio, this operation is a gradient-guided Metropolis-style acceptance rule rather than a general Hastings-corrected transition [9, 10].

Allowing occasional objective-increasing substitutions enables the optimizer to escape locally favorable sequence states that would trap a purely greedy search.

### S6.2 Temperature annealing

The acceptance temperature is geometrically annealed over the optimization trajectory. Given an initial temperature *T*_0_ and a half-life of *h* proposal steps,

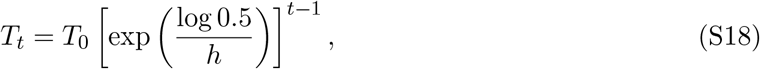

or equivalently,

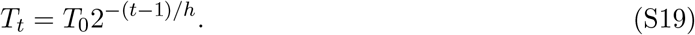

The default RF3 cycling configuration uses

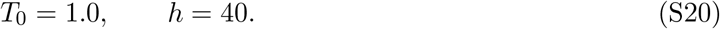

The resulting simulated-annealing schedule favors exploration during early proposal steps and progressively shifts toward local refinement [11].

### S6.3 Gradient recomputation following acceptance

When a mutation is accepted, RFO recomputes a new RF3 gradient at the accepted sequence. The next proposal therefore reflects the local objective geometry around the new sequence rather than continuing to use a gradient evaluated at the preceding state.

If a proposed mutation is rejected, the current sequence and its previously computed gradient are retained.

### S6.4 Optional multi-checkpoint consensus optimization

The implementation can load multiple RF3 checkpoints. Three strategies are implemented for constructing the mutation gradient:

i. use the first checkpoint;
ii. randomly select one checkpoint at each proposal step; or
iii. compute normalized gradients from all checkpoints and average them.

When several checkpoints are present, each model independently evaluates the candidate sequence. The code supports either a majority vote over temperature-dependent model-specific Metropolis decisions or a deterministic majority vote based on whether each model’s objective improves.

These ensemble modes are optional and are distinct from the RF3–Boltz cross-model cycling strategy used by the main RFO protocol.

### S6.5 Recorded optimization trajectory

Every evaluated GradMCMC proposal is recorded together with its sequence, objective value, acceptance state, and auxiliary model metrics.

Predicted RF3 structures associated with evaluated proposals are written to a separate trajectory directory. When the RF3 optimization branch is invoked by the outer cycling driver, the driver identifies the history entry with the lowest stored proposal loss and transfers the corresponding predicted structure to the subsequent inverse-folding stage.

The public cycling configuration performs five internal GradMCMC proposal steps per RF3 branch invocation.

## S7 Optimization objectives

RFO exposes differentiable objectives derived from RF3 confidence predictions and RF3 distogram outputs. All optimization terms are represented as losses to be minimized.

For the binder-design benchmarks described in the main text, mean interface PAE is used as the principal gradient objective. Geometry-sensitive design tasks can instead use composite confidence and distance objectives.

## S7.1 Mean interface predicted aligned error

Let *E_ij_* denote the expected predicted aligned error between tokens *i* and *j*, obtained by converting the RF3 PAE probability distribution into an expected error.

Let

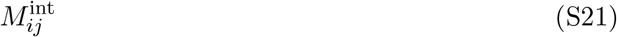

denote the inter-chain pair mask used by the RF3 metric implementation.

The mean interface PAE objective is

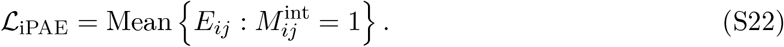

Lower values correspond to greater predicted confidence in the relative placement of interacting molecular components.

### S7.2 Global PAE and PDE

RFO additionally exposes global mean PAE,

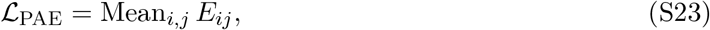

and mean predicted distance error (PDE) derived from the corresponding RF3 confidence head.

### S7.3 pLDDT objective

RF3 predicts binned pLDDT values for atoms. Expected pLDDT values are computed from the probability distribution, restricted to real heavy atoms, averaged within residues, and finally averaged over residues.

Because RFO minimizes all objective functions, confidence maximization is implemented as

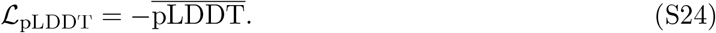

### S7.4 Distogram representation

Let

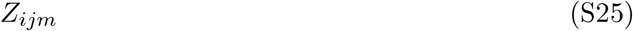

denote the RF3 distogram logit for token pair (*i, j*) and distance bin *m*. The normalized distribution is

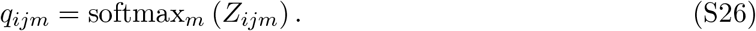

The implementation associates distogram bins with distances spanning approximately

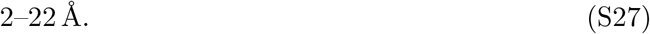

If *d_m_* denotes the representative distance associated with bin *m*, the expected token–token distance is

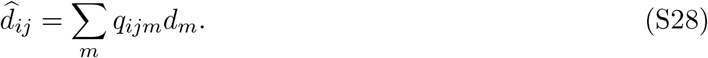

This representation provides a differentiable structural proxy without requiring optimization directly through sampled Cartesian coordinates.

### S7.5 Inter-chain contact loss

For a contact cutoff *d_c_*, RFO constructs a distance distribution restricted to bins below the cutoff,

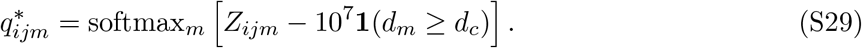

The corresponding pairwise contact loss is

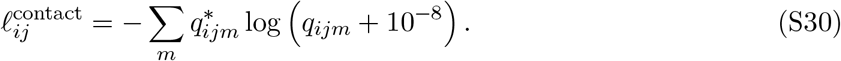

For the inter-chain objective, the first chain is treated as the binder and all remaining chains are treated as target context.

For every binder residue, RFO selects the *k* lowest binder–target pair losses. The default inter-chain parameters are

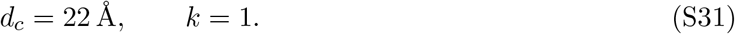

The residue-level contact terms are subsequently sorted and averaged.

### S7.6 Intra-chain contact loss

The intra-chain contact objective is computed analogously between residues of the binder chain.

Residue pairs separated by fewer than nine positions in sequence are excluded. For each binder residue, the two lowest valid contact losses are retained.

The default parameters are

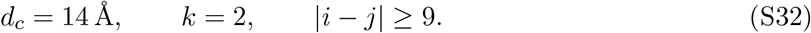

The combined contact objective is

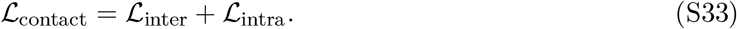

### S7.7 Interface distogram entropy

The entropy of the predicted distance distribution for token pair (*i, j*) is

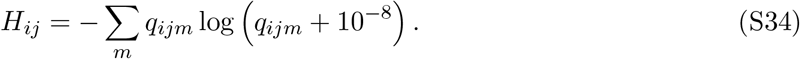

The interface-entropy objective averages this quantity over cross-chain token pairs,

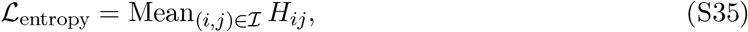

where *ℐ* denotes the set of binder–target token pairs. A proximity-weighted variant assigns the weight

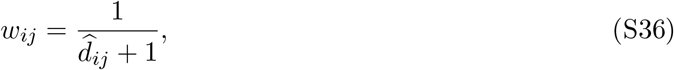

giving

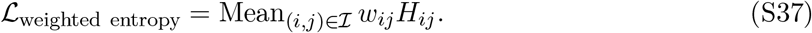

This objective preferentially reduces distance uncertainty for molecular pairs that are already predicted to be spatially proximal.

### S7.8 Distogram cross-entropy

RFO additionally implements a distogram cross-entropy objective using token-level distances derived from the supplied structure as the reference.

This objective is useful when sequence optimization should reinforce or preserve an existing predicted geometry rather than exclusively optimize generic confidence or contact formation.

### S7.9 Distance and geometric constraints

For geometry-sensitive optimization, the codebase contains a differentiable interval-based distance loss.

For predicted distance *d* and desired interval [*d*_min_, *d*_max_], the penalty is

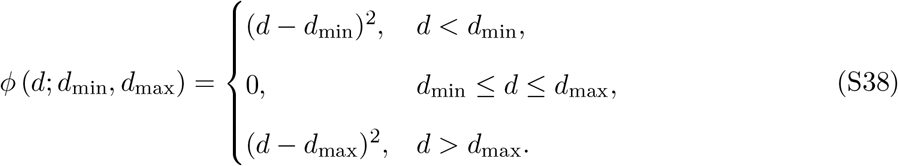

For *K* geometric constraints, the composite loss is

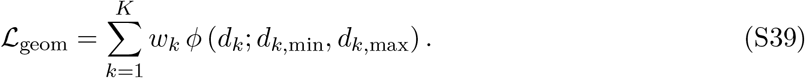

The same composite-loss framework supports pLDDT constraints at the chain or motif level. Thus, enzyme optimization can simultaneously constrain the relative placement of catalytic residues while encouraging a confidently predicted surrounding scaffold.

The repository also contains a newer `DistanceConstraint` utility that extracts expected distogram distances between specified residue pairs and records them as metrics. This utility itself returns zero optimization loss. Differentiable geometric optimization is provided by the interval-penalty formulation described above.

### S7.10 iPTM

The RF3 wrapper exposes iPTM as a confidence metric. In the current implementation, the internal iPTM calculation is converted to a Python scalar before being returned to the optimization interface and is therefore detached from the autograd graph.

Consequently, the benchmark gradient optimization described here does not rely on iPTM as a differentiable objective. iPTM is instead used as an evaluation and filtering metric. Mean interface

PAE is the principal differentiable confidence objective used for the binder benchmarks.

## S8 Structure-prediction and inverse-folding cycling

### S8.1 Stochastic branch selection

The outer RFO driver performs *C* completed optimization cycles.

At cycle *c*, a random number

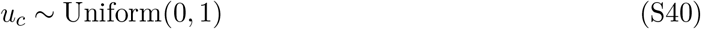

is sampled.

The RF3 gradient branch is selected when

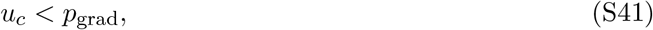

and the Boltz branch is selected otherwise.

The default configuration uses

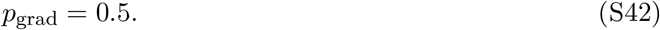

Thus, over a sufficiently long trajectory, gradient-derived sequence optimization and independent structure-prediction cycling are sampled with equal probability.

A successful structure-prediction branch is always followed by MPNN redesign.

If either the structure-prediction step or MPNN redesign fails, the cycle is retried and is not counted as a completed optimization cycle. For *C* requested cycles, the implementation permits at most

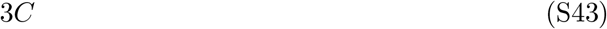

branch attempts.

### S8.2 Boltz structure-prediction branch

When the Boltz branch is selected, RFO converts the current complex into an AF3-style molecular specification and subsequently into the YAML format used by Boltz.

The current sequence of chain A is refreshed from the most recent RFO structure while target and ligand specifications are preserved.

When an explicit AF3-style template is supplied, RFO copies the template and updates only the chain-A sequence. This allows metadata that are not recoverable from a PDB structure, such as a SMILES ligand representation or embedded MSA specification, to remain constant throughout optimization.

Boltz is then run as a frozen structure predictor.

The repository defaults are

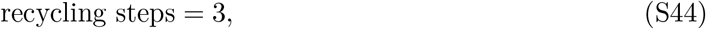

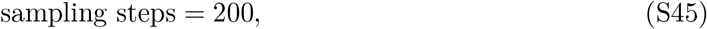

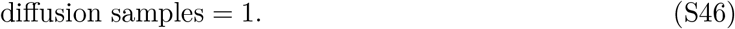

The predicted model-0 structure is used for the subsequent inverse-folding stage.

These quantities describe the software defaults. Experiment-specific production configurations, where different, supersede the defaults.

### S8.3 High-confidence residues fixed after Boltz prediction

Following Boltz prediction, RFO identifies binder residues that should be preserved during MPNN redesign.

The Boltz output stores atom-level confidence values in the mmCIF B-factor field. For each residue of chain A, RFO computes the mean confidence over its atoms.

A residue is classified as high confidence when

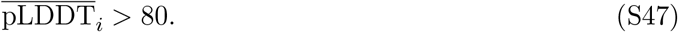

To avoid fixing isolated high-confidence residues, only contiguous high-confidence stretches containing at least five residues are retained.

Residues in these stretches are held fixed during the immediately following MPNN redesign.

### S8.4 Interface residues fixed after RF3 optimization

Following an RF3 gradient branch, RFO identifies chain-A residues that contact chain B. A binder residue is classified as an interface residue when any of its atoms lies within

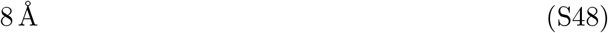

of an atom in chain B.

These residues are fixed during the subsequent MPNN redesign.

This 8 Å MPNN-fixation cutoff is distinct from the optional 5 Å interface definition used to restrict positions accessible to the gradient mutation operator.

### S8.5 ProteinMPNN and LigandMPNN redesign

Following either structural branch, RFO performs structure-conditioned inverse folding.

ProteinMPNN is used by default for protein-only systems, whereas LigandMPNN can be used when ligand or other non-protein atomic context should condition the sequence-design step [6, 7].

Only residues from the designated design chain that are not in the branch-specific fixed set are passed as designable positions.

If *ℱ* denotes the fixed chain-A residue set, the inverse-folding design set is

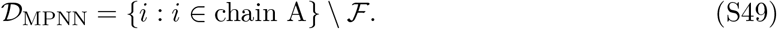

Target chains are implicitly fixed.

The default MPNN sampling temperature is

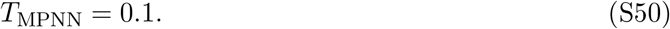

The public worker generates one sequence by default and uses the first design returned by the MPNN inference engine.

The redesigned structure is written to disk and becomes the complete input structure for the following RFO cycle.

## S9 Complete RFOptimization procedure

The complete optimization protocol can be summarized as follows.

**Algorithm S1, step 1:** Initialize the current state from an input biomolecular complex *X*^(0)^ and designate chain A as the sequence-design chain.

**Algorithm S1, step 2:** At outer optimization cycle *c*, sample the RF3 gradient branch with probability *p*_grad_ or the Boltz prediction branch with probability 1 − *p*_grad_.

**Algorithm S1, step 3:** If RF3 is selected, evaluate the current sequence, compute the objective gradient with respect to the residue-type input, generate discrete gradient-derived mutations, and accept or reject proposals using the temperature-dependent Metropolis rule.

**Algorithm S1, step 4:** Repeat the RF3 proposal procedure for the configured number of internal GradMCMC steps and retain the structure selected from the RF3 trajectory.

**Algorithm S1, step 5:** If Boltz is selected instead, predict the all-atom structure of the current sequence using the independently trained Boltz predictor.

**Algorithm S1, step 6:** Construct a branch-specific set of chain-A residues to preserve during inverse folding.

**Algorithm S1, step 7:** Redesign the remaining chain-A residues using ProteinMPNN or LigandMPNN.

**Algorithm S1, step 8:** Set the MPNN-redesigned complex as the input state *X*^(*c*+1)^.

**Algorithm S1, step 9:** Repeat until the requested number of successfully completed outer cycles has been obtained.

## S10 Default implementation parameters

Table S1 summarizes the principal defaults in the supplied RFO implementation.

These parameters document the public software behavior and should be distinguished from experiment-specific overrides used for individual production benchmarks.

## S11 Benchmark-specific computational protocols

### S11.1 Protein–protein binder optimization

RFO was evaluated on RFdiffusion-generated protein binders for 13 protein targets previously used in computational binder-design studies [1].

The starting complexes were treated as local sequence-optimization seeds rather than regenerated from scratch. Chain A was designated as the binder.

Target MSAs and structural templates were supplied to RF3 where appropriate, and mean interface PAE was used as the principal sequence-optimization objective.

Each starting design was followed for up to 10 outer RFO cycles.

The initial and optimized sequences were independently refolded using AlphaFold3 for held-out evaluation.

A design was classified as successful when the AF3 prediction simultaneously satisfied

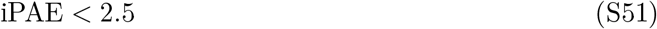

and

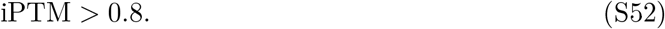

Importantly, these criteria are applied to the independent AF3 evaluation rather than to the RF3 objective used to construct sequence mutations.

### S11.2 Cyclic-peptide optimization

Cyclic-peptide starting structures were generated using RFpeptides for four targets:

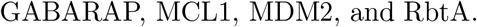

Chain A was marked as cyclic during structure prediction.

Mean interface PAE was used as the RF3 optimization objective.

Final success was evaluated using held-out AF3 refolding under the same criterion,

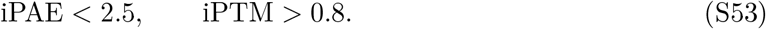

**S11.3 Small-molecule binder optimization**

Small-molecule binder optimization was evaluated for four ligands:

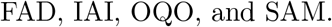

Ligands were retained as explicit all-atom molecular context throughout the structure-prediction stage.

When required, ligand identity was preserved through the AF3-style specification using a CCD identifier or SMILES representation.

Mean interface PAE was used as the RF3 sequence-optimization objective.

LigandMPNN can be used for inverse folding when ligand-conditioned protein sequence design is required.

Final success was evaluated using held-out AF3 refolding under the same iPAE/iPTM criterion.

### S11.4 Atomic-motif enzyme optimization

RFO was additionally evaluated on 24 targets from the atomic motif enzyme benchmark. Initial motif-scaffolded structures were used as local optimization seeds.

The enzyme objective combines predicted structural confidence with geometric constraints on catalytically important residues.

In general form,

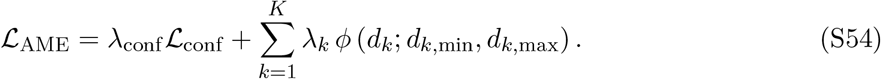

The confidence term promotes a confidently predicted scaffold, whereas the geometric terms penalize deviation from the required catalytic arrangement.

Because the geometric objective is computed from the all-atom model representation, constraints can be defined between selected positions within the catalytic motif while the surrounding scaffold remains free to change sequence.

## S12 Held-out and cross-model evaluation

### S12.1 AlphaFold3 as a held-out evaluator

AlphaFold3 [3] is not used to construct RF3 gradient proposals in the reported binder benchmark protocol.

Instead, it is used as an external structure-prediction evaluator.

This separation reduces the likelihood that an apparent improvement is solely the result of exploiting the confidence landscape of the model that provided the optimization gradient.

The same AF3 filter is applied to the starting and optimized sequences.

For trajectory-level analyses, a “best-of-*k*” pass criterion can be used: a starting design is counted as rescued if at least one evaluated sequence among the first *k* optimization cycles satisfies the final AF3 filters.

### S12.2 Three-model consensus evaluation

To evaluate whether optimized sequences transfer across independently trained predictors, designs are evaluated with RF3, Boltz, and AF3.

For an iPTM threshold *θ*, define the model-specific indicator

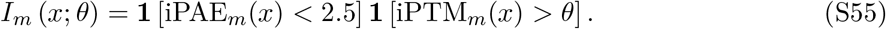

The three-model consensus criterion is

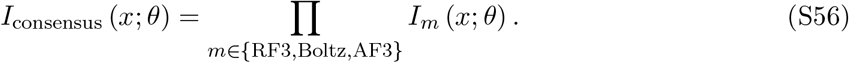

A sequence therefore passes only if all three predictors simultaneously satisfy both the iPAE and iPTM requirements.

The iPTM threshold is swept from

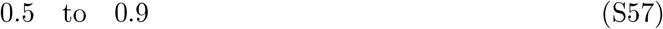

in the cross-model analysis.

The primary operating point reported in the main text is

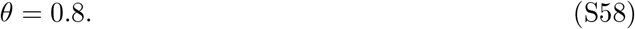

## S13 Ablations and external baselines

### S13.1 Cycling-only ablation

The cycling-only ablation removes gradient-guided RF3 sequence optimization by setting

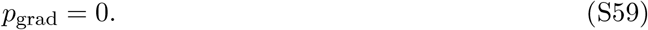

Every optimization cycle therefore consists of Boltz structure prediction followed by the same MPNN redesign procedure used in the full method.

Starting designs, number of cycles, and final evaluation criteria are otherwise kept matched to the full RFO protocol.

This ablation isolates the contribution of gradient-derived objective information from the contribution of repeated structure-conditioned inverse folding alone.

### S13.2 Protein Hunter and BindCraft

Protein Hunter is used as an independently developed cycling-based baseline. BindCraft provides a predictor-guided hallucination baseline [8].

Where method input requirements permit, the resulting designs are evaluated using the same downstream structure-prediction criteria to enable a controlled comparison of filter-passing efficiency.

### S13.3 Computational-efficiency calculation

For computational-efficiency analyses, the amortized GPU cost per filter-passing design is calculated as

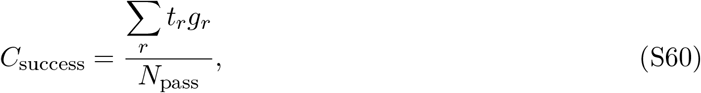

where *t_r_* is the wall-clock duration of run *r, g_r_* is the number of GPUs used by that run, and *N*_pass_ is the number of resulting sequences that satisfy the common final filter.

Failed and non-passing trajectories remain part of the numerator because they represent real computational expenditure required to obtain a passing candidate.

## S14 Software execution and reproducibility

RFO is implemented in Python.

The host-side cycling driver performs lightweight structure and sequence bookkeeping and launches RF3, Boltz, and MPNN inference using GPU-enabled Apptainer containers.

Container locations, model checkpoints, caches, and external molecular-data mirrors are configurable through environment variables. This allows the same Python optimization logic to be executed against different installations without hard-coding model assets into the optimization procedure.

The public command-line interface accepts a YAML configuration file. Parameter precedence is

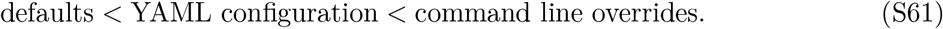

At minimum, the user specifies

i. an initial biomolecular structure, and
ii. an output directory.

Optional parameters include the RF3/Boltz routing probability, number of cycles, optimization objective, chain-specific MSAs, target templates, Boltz sampling parameters, MPNN temperature, cyclic-chain handling, and random seed.

For each completed cycle, RFO retains the predicted structure, MPNN output, and model-specific metrics.

The RF3 branch additionally records the sequence and loss associated with individual GradMCMC proposals and stores predicted structures generated during the RF3 search trajectory.

At the end of a run, RFO writes a JSON record containing the completed-cycle trajectory, selected branch at each cycle, MPNN-designed sequence, model metrics, and wall-clock duration.

Runtime measurements reported in the main text were obtained using NVIDIA A100-class GPUs. For exact reproduction of a benchmark run, the following information should be archived together:

i. the initial structure;
ii. the YAML run configuration;
iii. RF3, Boltz, and MPNN checkpoint identifiers;
iv. target MSA and structural-template inputs;
v. ligand CCD or SMILES specification where applicable;
vi. random seed; and
vii. the exact RFO software commit.

## S15 Implementation notes relevant to reproducibility

The RFO repository contains both the current cycling implementation and earlier experimental loss and optimization components.

For reproducibility, it is useful to distinguish three classes of functionality.

First, the current outer cycling driver directly uses the RF3 GradMCMC branch, Boltz structure prediction, and MPNN redesign.

Second, the codebase contains differentiable specialized objective classes, including interval-based distance constraints and motif-level confidence constraints, which are useful for geometry-sensitive optimization tasks.

Third, some quantities are exposed as model metrics but are not currently gradient-bearing objectives. In particular, the current iPTM implementation converts the calculated score into a Python scalar before returning it to the optimizer.

This distinction ensures that the mathematical description of an optimization objective corresponds to the actual gradient path used in the implementation, rather than simply to the presence of a metric in the software interface.

---

**Table S1:** Principal default parameters in the RFOptimization cycling implementation.

| Parameter | Default | Role |
| --- | --- | --- |
| Gradient-branch probability | 0.5 | Probability that a cycle begins with RF3 rather than Boltz |
| Total completed cycles | 10 | Number of outer RFO cycles |
| Random seed | 42 | Host-side routing and cycle-specific MPNN seeding |
| Boltz diffusion samples | 1 | Structural samples generated per Boltz invocation |
| Boltz recycling steps | 3 | Boltz recycling iterations |
| Boltz sampling steps | 200 | Boltz structure-sampling iterations |
| RF3 recycles | 10 | RF3 recycles used during gradient optimization |
| RF3 diffusion batch size | 1 | Number of RF3 structure samples |
| GradMCMC steps per RF3 branch | 5 | Internal sequence-proposal steps in the default cycling configuration |
| Maximum substitutions per proposal | 1 | Maximum number of amino-acid changes retained in one GradMCMC proposal |
| Initial MCMC temperature | 1.0 | Initial Metropolis acceptance temperature |
| Temperature half-life | 40 steps | Geometric annealing half-life |
| MPNN sequences per redesign | 1 | Default inverse-folding batch size |
| MPNN sampling temperature | 0.1 | Sequence-sampling temperature |
| Boltz confidence threshold | 80 | Mean residue confidence required for fixation after Boltz prediction |
| Minimum high-confidence run | 5 residues | Minimum contiguous segment fixed after Boltz prediction |
| RF3-to-MPNN interface cutoff | 8 Å | Contact cutoff for fixing binder residues after an RF3 branch |
| Gradient interface-only cutoff | 5 Å | Optional cutoff defining residues accessible to gradient mutation |
| Maximum cycle attempts | 3*C* | Retry budget for *C* requested completed cycles |

